# The role of tectonics and hydrothermalism in early human evolution at Olduvai Gorge

**DOI:** 10.1101/632414

**Authors:** Ainara Sistiaga, Fatima Husain, David Uribelarrea, David Martín-Perea, Manuel Domínguez-Rodrigo, Troy Ferland, Katherine H. Freeman, Fernando Diez-Martín, Enrique Baquedano, Audax Mabulla, Roger E. Summons

## Abstract

Hominin encephalization has been at the centre of debates concerning human evolution with a consensus on a greater role for improved dietary quality. To sustain the energetic demands of larger brains, cooking was likely essential for increasing the digestibility and energy gain of meat and readily available, yet toxic starches. Here, we present the oldest geochemical evidence for a landscape influenced by tectonic activity and hydrothermal features that potentially shaped early hominin behaviour at Olduvai Gorge. Although use of fire at this time is controversial, hot springs may have provided an alternative way to thermally process dietary resources available in the 1.7 Myo Olduvai wetland. Our data supports the presence of an aquatic-dominated landscape with hydrothermal features that offered hominins new opportunities to hunt and cook readily available tubers and herbivore prey at the emergence of the Acheulean technology. Future studies should further examine whether hydrothermalism similarly influenced other critical aspects of human evolution.

## Introduction

Reconstruction of specific paleoenvironments within the cradle of humanity may yield evolutionary insights that illuminate the history of the human species. Many environmental factors including aridity^1,2^, climate variability^3^, topographic roughness^4^, volcanoes^5^, and taphonomic bias^6^ have been suggested as drivers of early human evolution. Olduvai Gorge, located in the Great Rift Valley in Tanzania, has yielded some of the most important hominin fossils in the world. Previous landscape reconstructions associated with early human evolution suggest that climate and vegetation at Olduvai was strongly correlated with precession-driven wet-dry cycles^7–10^ imposed on a general aridification trend^11,12^. The 100,000-year precessional cycles caused the basin vegetation to vary between closed forested ecosystems during wetter periods^7,13^, and C_4_ grass-dominated environments during arid periods.

Debates about the emergence, co-existence, and evolution of the Acheulian have been revitalized by the discovery of FLK West (FLK-W) in Bed II of Olduvai Gorge, Lower Augitic Sandstones (LAS), 1.7 Myo^14–16^, which contains the oldest examples of Acheulean technology, including symmetric bifacial stone hand-axes^15^. This site is contemporaneous with the HWK sites, whose abundant lithic assemblages have been classified as Oldowan, containing simple core-flaked tools^17,18^. The simultaneous presence of Acheulian and Oldowan tools suggests that both technologies co-existed on the same landscape and could have had different uses by early humans^19,14^. Uribelarrea and colleagues’ reconstructions^14^ of the LAS landscape show that the Oldowan occurrences at HWK sites are part of a proximal shallow braided stream system, whereas FLK-W occupies a more distal location towards the lake floodplain, in which the braided streams had coalesced into a wider channel (Fig.1). Within this interpretation, vegetation at HWK locations may have been distributed less densely, whereas vegetation at FLK-W may have been denser, creating a canopy cover, and may have been wetter. Open-vegetation fauna at HWK has been recently documented^20^, while more mixed-habitat fauna has been found at FLK-W. Here we present paleolandscape reconstructions for Bed II, LAS, 1.7 Myo, which corresponds with the appearance of the oldest Acheulean^14–16^, using high-resolution lipid biomarker and isotopic signatures.

**Figure 1:**
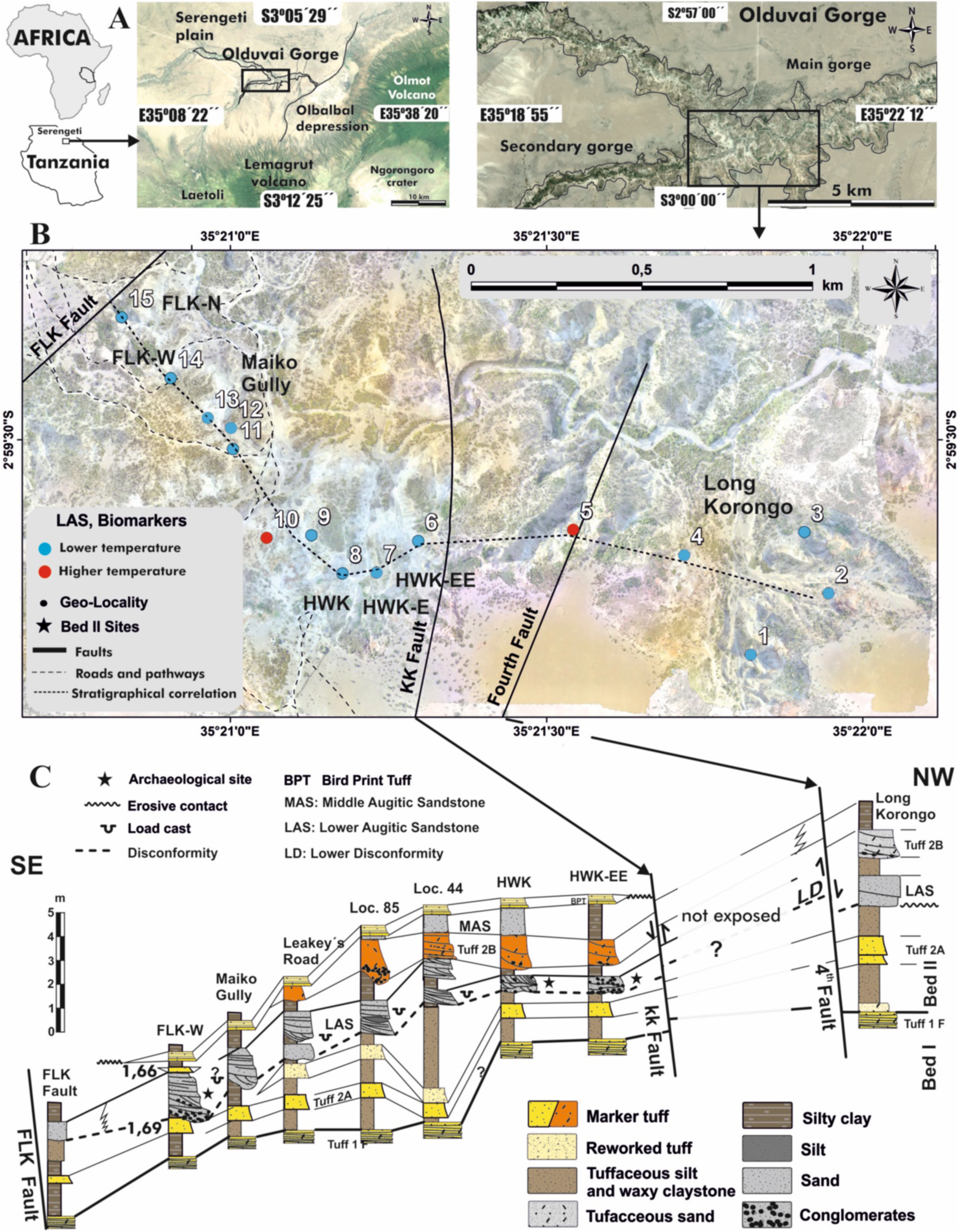
(A) Geographical location of Olduvai Gorge and the study area. (B) Detailed map of the study area, based on the orthoimage obtained with unmanned aerial vehicle. The blue and red dots indicate the position of the samples described in this study. The red colour indicates the presence of biomarkers indicative of high temperature hydrothermal activity.

The discovery of Acheulean technology at FLK-W suggests that complex cognition was present 1.7 Myo at Olduvai Gorge^15^. The metabolic demands of larger brains, together with small guts and weakened dentition, point to increased food processing, which in the absence of controlled fire could have included grinding to increase digestibility^21^.

Our analysis reveals a wetland ecosystem with groundwater-fed rivers and aquatic plants, sourced by hydrothermal features. Hydrothermal activity may have influenced the use of the space at Olduvai Gorge, as hot springs may have enabled hominins to cook animal tissues and tubers with minimal effort. The possibility of using such hydrothermal features for thermal processing has not yet been suggested, and may support the presence of hydrothermal cooking during a pre-fire stage in human evolution.

## Geologic setting

In the study area, the base of Bed II is a waxy claystone corresponding to a lake margin of very low energy. Tectonic reactivation of the region caused descent of a block limited by the FLK and 5^th^ faults, creating a large graben between them (Fig.1). The resultant decrease in the regional base level favoured incision of the entire drainage network that emptied into the central lake (FLK-5^th^ fault block). The resulting paleosurface is known as Lower Disconformity (LD) and is found across almost the entire gorge. Between HWK-EE and FLK-W, the same river system that formed the LD, deposited so-called Lower Augitic Sandstones (LAS), dated at 1.7 Myr ago^15^. The continuity of this fluvial deposit overlying the LD in the whole study area^14,22,23^, has allowed it to be used as a chronostratigraphic unit, its base being an isochronous paleosurface. For this type of work to be sound, centimetre-scale control of the stratigraphy in the field is required and this has enabled the LD paleosurface to be located within a 100 meters thick record and despite the study area being affected by the 4^th^, KK and FLK faults with corresponding drag folds. Thus, it has been the geological work in LAS conducted since the discovery of FLK-W in 2012 that enabled the precise location of these 15 samples in the corresponding isochrone.

The contemporaneous presence of FLK-W and LAS support the notion that Acheulian and Oldowan technologies coexisted, reinforcing the cladogenetic interpretation of the emergence of the Acheulian and its typological co-existence with other industries. Other authors, Stanistreet^24^ have also documented the presence of LAS in FLK (Loc.45), according to Hay’s^22^ geological interpretation, although they later changed their interpretation to suggest the presence of FLK-W in a more modern augitic unit (MAS), after the Tuff IIB and therefore not contemporary with the HWK deposits^25,26^, which would lead to interpreting technology following an anagenetic evolution (SI1).

## Results and discussion

Beginning ~3 Myr ago, pronounced changes in East Africa due to progressive uplift and rifting allowed the gradual replacement of C_3_ vegetation with C_4_-dominated grasslands as shown by plant biomarker and isotopic data^27^. Previous paleoenvironmental reconstructions point to lake expansion-retraction cycles, associated with wetter precession-driven lake refreshing spikes between 1.9 and 1.8 Myo^9,12,28^, followed by a subsequent major shift towards a drier environment^11,12,27,29^. At approximately 1.7 Myo, despite the overarching drying trend, tectonic activity enabled a general reactivation of the drainage network and a progradation of the river systems at Olduvai Gorge. This created wetter intervals superimposed on an overall long-term aridification trend ^9,29,30^.

Plant biomarkers, especially epicuticular *n*-alkane leaf waxes, have been widely used to reconstruct paleoclimate and paleovegetation changes in ancient environments^31,32^. All LAS samples analysed yielded significant amounts of *n*-alkanes. Most of the samples present an *n*-alkane distribution dominated by the long-chain, odd-numbered C_29_, C_31_, and C_33_ *n*-alkanes typical of terrigenous inputs^31,33^, while mid-chain length C_23_ and C_25_ *n*-alkanes, which have been attributed to the presence of aquatic macrophytes, are also present in significant proportions^34,35^.

While aquatic macrophytes currently thrive in most high-altitude African lakes, their historical presence at the Olduvai Gorge paleolake was initially confirmed in Bed I^35^. The biomolecular proxy *P*_aq_ expresses the proportion of submerged and floating macrophytes versus emergent macrophytes and terrestrial plants^34^. Most LAS samples have *P*_aq_ values between 0.2-0.4 (Fig.2E), which correspond to the values suggested for emergent macrophytes, such as plants from the genus *Typha*, fossil imprints for which have been found at Olduvai Gorge within Lower Bed II^36^. *P*_aq_ values below 0.1 are indicative of greater proportions of terrestrial input, confirming that the LAS environments sampled are dominated by emergent aquatic plants, similar to the wetlands present today in Africa^34^. Intermediate values between 0.1 and 0.4 may reflect a mixed input, pointing to a similar setting like that described for Lake Rutundu in Kenya^34^. Here, abundant floating aquatic plants such as *Potamogeton* in shallow waters with a fringe of emergent macrophytes are surrounded by ericaceous shrubland.

**Figure 2:**
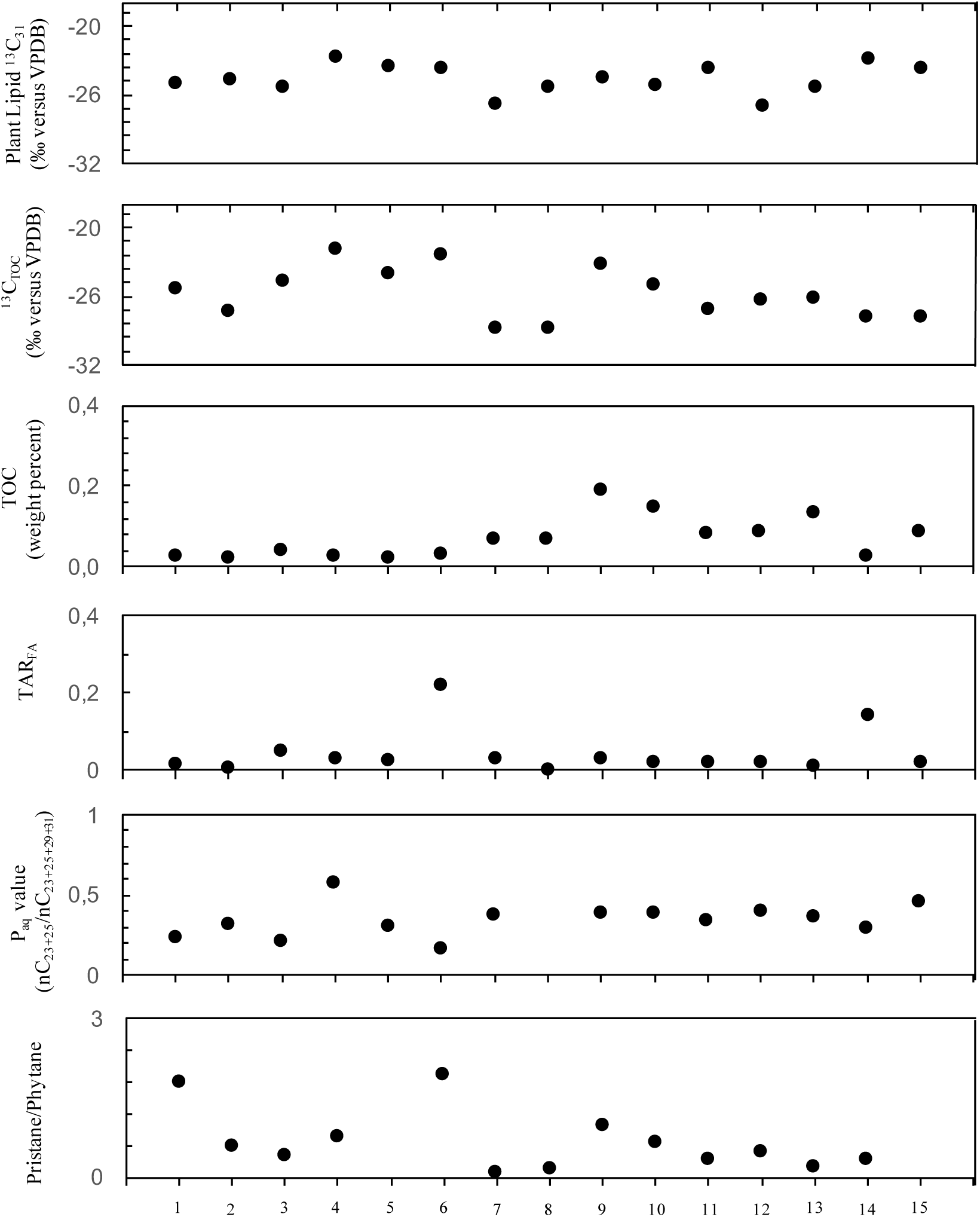
Plot of various organic geochemical proxies for organic matter preserved in outcrops from Eastern fluvial lacustrine deposits (eastern lake-margin) dated up to 1.7 Myr ago at Olduvai Gorge. From top to bottom: (A) Plant lipid δ^13^C values for *n*-C_31_ alkane (higher values, more C4 vegetation); (B) Carbon isotopic composition of total organic carbon (higher values, more C4 vegetation); (C) Total organic carbon, weight percent (higher values, more organic matter production); (D) Terrigenous to aquatic n-alkanoic acids ratio reflecting the importance of terrigenous and aquatic sources (C_24_ + C_26_ + C_28_)/(C_14_ + C_16_ + C_18_) (higher values, more terrestrial input); (E) Ratios of macrophytic lipids (*n*-C_23_ + *n*-C_25_) relative to macrophytic and terrestrial lipids (*n*-C_23_ + *n*-C_25_ + *n*-C_29_ + *n*-C_31_) (higher values → more macrophytic input); (F) Ratios of pristane to phytane (values above 2 often indicative of more terrestrial plant input, values below 1 indicative of anoxic conditions, values around 1 indicative of oxic/anoxic alternating conditions).

Only LAS 5 falls above 0.4 with a *P*_aq_ value of 0.56, which is consistent with a floating/submerged macrophyte signal. LAS 6 from the HWK-NE archaeological site produced the lowest *P*_aq_ value of 0.16, which is very close to the terrestrial plant threshold, (>0.1) suggesting an increased terrestrial input in this area.

The pristane-to-phytane ratio is a hydrocarbon-based proxy used to assess the redox state of sedimentary environments. Values greater than 2 suggest overall oxic conditions, values less than 1 indicate persistently anoxic conditions, and values between indicating alternating oxic-anoxic conditions^33^. Two samples, LAS 1 and 6, have pristane-to-phytane ratios of 1.68 and 1.94, respectively, suggesting that they originate from intermittently oxic environments. Other LAS samples have values between 1 and 0, suggesting aquatic settings not permanently flooded but not desiccated either (Fig.2F). Similarly, the terrigenous-to-aquatic fatty acid ratio (Fig.2D), which reflects the proportion of terrestrial to aquatic sources^37^, shows more elevated values for both LAS 6 and LAS 14, which suggests increased terrestrial plant inputs in both HWK-NE and FLK-W. Accordingly, TOC values were low in most LAS samples, with nitrogen levels below detection limits (Fig.2C).

Although *n*-alkanes have little chemotaxonomic significance for temperate plants^32,38^, recent work by Bush and McInerney^32^ showed that savanna plants are more accurately identified based on their carbon chain length distributions that their temperate counterparts. Discriminant analyses using C_29_, C_27_, and C_31_ distributions; C_29_, C_27_, and C_33_ distributions, and C_27_, C_29_ and C_35_ distributions compared with previously published data^32,38^, suggest that the terrestrial *n*-alkanes from our samples might have a savanna angiosperm or C_4_ graminoid origin (Fig.3A,B,C). Likewise, distributions of C_27_, C_29_ and C_23_ *n*-alkanes in most LAS samples compared to data collected by Bush and McInerney^32^ point to an angiosperm origin, except for LAS 14 from FLK-W, which has a major C_23_ *n*-alkane contribution, meaning it potentially originated from the genus *Sphagnum*, whose mosses require cover resulting in low-light, humid conditions (Fig.3D).

**Figure 3:**
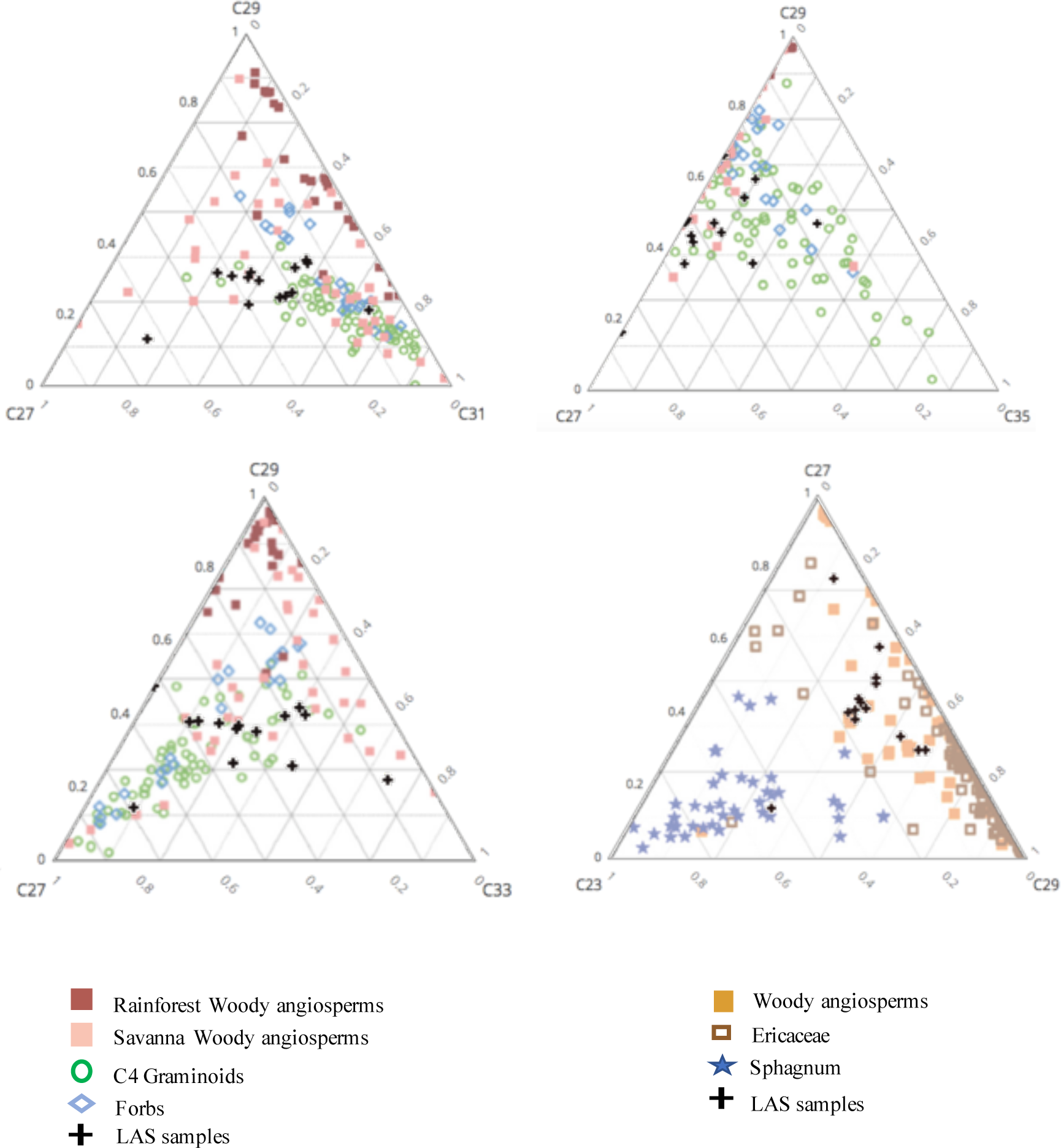
Ternary diagrams of the n-alkane chain-length abundances of the LAS samples compared to African rainforest woody and savanna angiosperms (A,B,C) from Bush and McInerney (2013) and Vogts et al., (2009); and temperate woody angiosperms, Ericaceae and sphagnum (D) from Bush and McInerney (2013).

δ^13^C values for the *n*-alkanes might provide a more insight into the dominant vegetation than δ^13^C_TOC_, which can carry signals from older and recalcitrant carbon phases. LAS samples have δ^13^C values that fall within wooded grassland ecosystem of African plant communities defined by UNESCO (Fig.2A). δ^13^C_31n_ values from LAS samples are within the suggested range for C_3_ graminoids^33^, but they may also point to the presence of aquatic macrophytes, which mostly use a C_3_ photosynthetic pathway^34,39,40^. Emergent plants from the genera *Typha, Cyperus, Hydrilla*, and *Potamogeton*, are often predominant in East African wetlands, and may display a similar range of δ^13^C values^34,40–43^. However, it is particularly difficult to specifically identify species given the wide range of δ^13^C for aquatic plants, which often depend on the differences in isotopic compositions of the source of carbon, water flow, turbulence, and individual physiological properties, such as the relative distribution of the plant above and below the water since this affects the plant’s access to HCO_3_^−^ or CO_2_^40^. Overall, our biomarker proxies point to a freshwater-fed wetland dominated by C_3_-grasses/aquatic plants and woody angiosperms, with high proportion of aquatic organic matter. This would be in accordance with previous geological and sedimentological descriptions that suggest a marsh-like depositional environment^22,44^.

Sedimentary δ^13^C_31n-alkane_ values range from −27.0‰ to −22.8‰, with an average value of −24.5‰ (SI2.A). Total organic carbon δ^13^C values (δ^13^C_TOC_) also show a small isotopic range of about 7‰, but correlate weakly with the δ^13^C_31_ values (r^2^ = 0.16). The δ^13^C_TOC_ is on average 1.2‰ more depleted than the δ^13^C_31_. These negative values, more consistent with an ecosystem with greater proportions of C_3_-like plants, might reflect the transport by fluvial waters of a non-soluble carbon pool eroded from older rocks, transported from upstream, or of microbial origin. Such depleted values might also reflect the effect of groundwaters, charged with CO_2_ with more negative values, that have passed through.

The analysis of the polar lipid fraction in our samples illuminates this phenomenon. Samples collected from the LAS layer show a very unusual distribution of functionalized lipids, including monoalkyl glycerol monoethers (MAGEs), which have been often interpreted as biomarkers for sulphate-reducing bacteria (SRB) and/or thermophilic organisms^45–47^, among other possible biological sources^48–50^. These glycerol ether lipids are often present in high concentrations at marine hydrothermal vents and cold seeps^51^, and in terrestrial geothermal sediments^46,47,52^. MAGEs are the predominant compound class in our alcohol fraction, constituting more than 80% of some of the samples (Fig.4D). A suite of MAGEs ranging from C_15_ to C_19_ with straight, branched, and unsaturated hydrocarbon chains dominated by the *n*-C_16_ MAGE followed by me-C_17:0_, which has been linked in the past to the activity of SRB, more specifically from the genus *Desulfobacter* or *Actinomycetes*^49,53^. MAGE distributions can be visualized in GC-MS analyses based on their predominant 205 Da fragment ion. Individual MAGE homologues and isomers were identified using elution order and full scan mass spectra (SI2.B)

**Figure 4:**
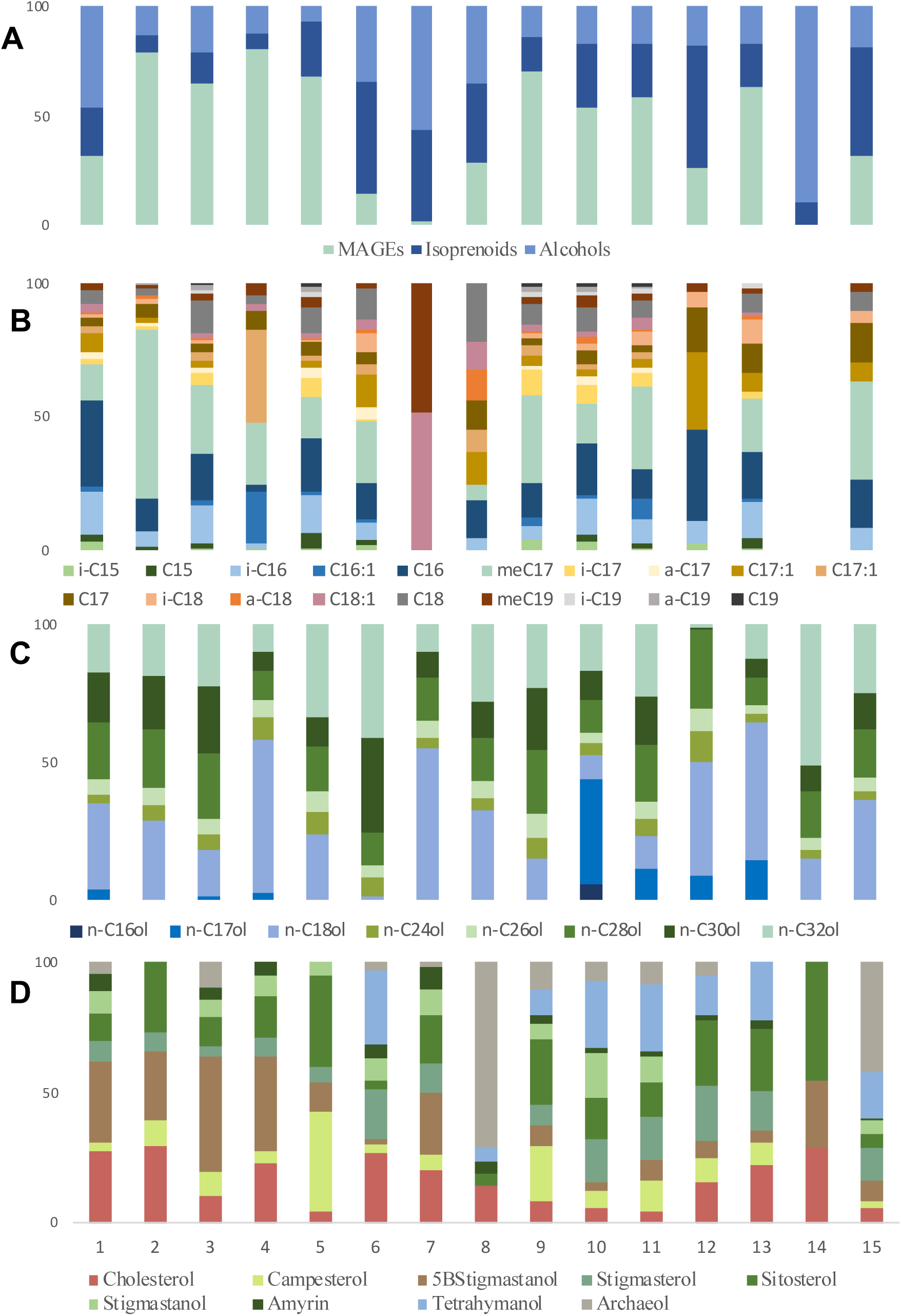
Histograms depicting the lipids identified in the GC-MS analysis of the polar fraction. Percentage distributions of (A) distribution of MAGEs, isoprenoids and *n*-alkanols within each sample, (B) monoalkyl glycerol monoethers, (C) *n*-alkanols, and (D) isoprenoids.

MAGEs identified in soils from non-SRB origin are often dominated by i-C_15_ and *n*-C_15_^49,50^ which are present in very low concentrations in our samples. Although our samples are mostly dominated by *n*C_16:0_ and Me-C_17:0_, LAS 1, 3-4, 7-8, and 12-13, contain significant amounts of MAGEs with a carbon chain length >17. Longer chain lengths, especially C_18:0_ and C_20:0_ have been associated with thermophilic bacteria of the genera *Aquificales*^46^. In particular, Jahnke and colleagues^46^ found that among the *Aquificales* bacteria, *Thermocrinis ruber* had a very distinctive lipid signature dominated by C_18_ MAGEs along with large amounts of C_20:1_, C_22:1_ and a C_21_ cyclopropane fatty acid (cy-C_21_) and where glycerol dialkyl ethers were subordinate. Strikingly, none of our samples yielded detectable glycerol dialkyl ethers, which are otherwise common in lake sediments, soils, and marine sediments. LAS 1-3, 7 and 11-13, all contained the C_20:1_, C_22:1_ fatty acids, which are the predominant fatty acids in *Thermocrinis ruber*^*46*^. Further, cy-C_21_ was tentatively identified in samples 5 and 10(Fig 5, SI2.D). *T. ruber* is a gram-negative hyperthermophilic bacterium that forms a separate lineage within *Aquificales*^*46*^, and was identified for the first time in streamer biofilm communities in Yellowstone hot spring outflow channels^54^. *T. ruber* grows optimally between 85°C and 95°C in waters with low salinity and near neutral pH. *T. ruber* is autotrophic, and has the ability to oxidize hydrogen, sulphur, and thiosulphate in the presence of oxygen, like the previously identified thermophilic bacteria *Aquifex* and *Hydrogenobacter*^*51,55*^. The lipid distributions identified within the LAS samples have been reported in few settings other than the Lower Geyser Basin of Yellowstone National Park, typified by features such as Octopus Spring and Bison Pool^46,52,56^ and Orakei Korako Spring^47^ in New Zealand.

**Figure 5:**
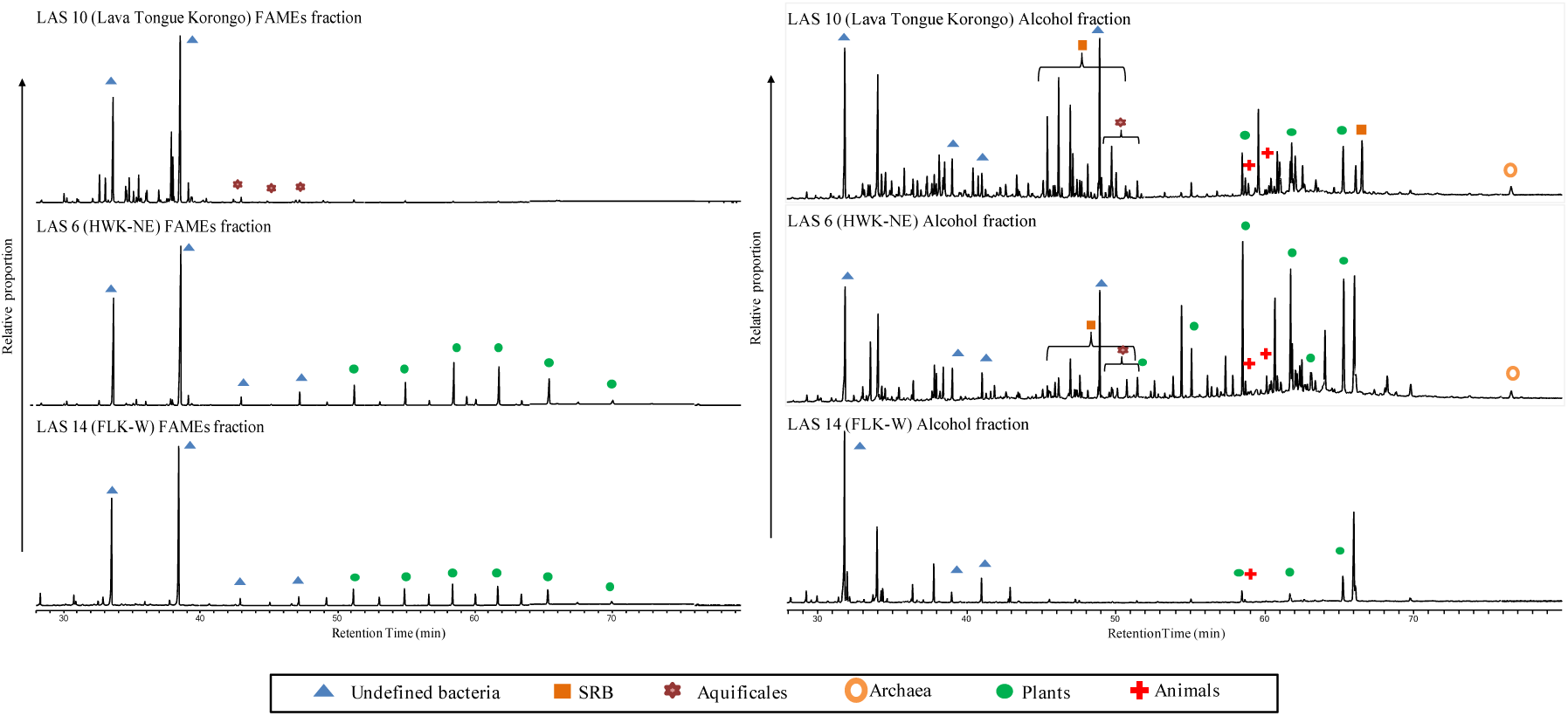
Partial total ion chromatogram from gas chromatography-mass spectrometry (GC-MS) analyses of fatty acids methyl ester (FAMEs) from samples 13,6,14 (right), and isoprenoids from the alcohol fraction (right).

**Figure 6:**
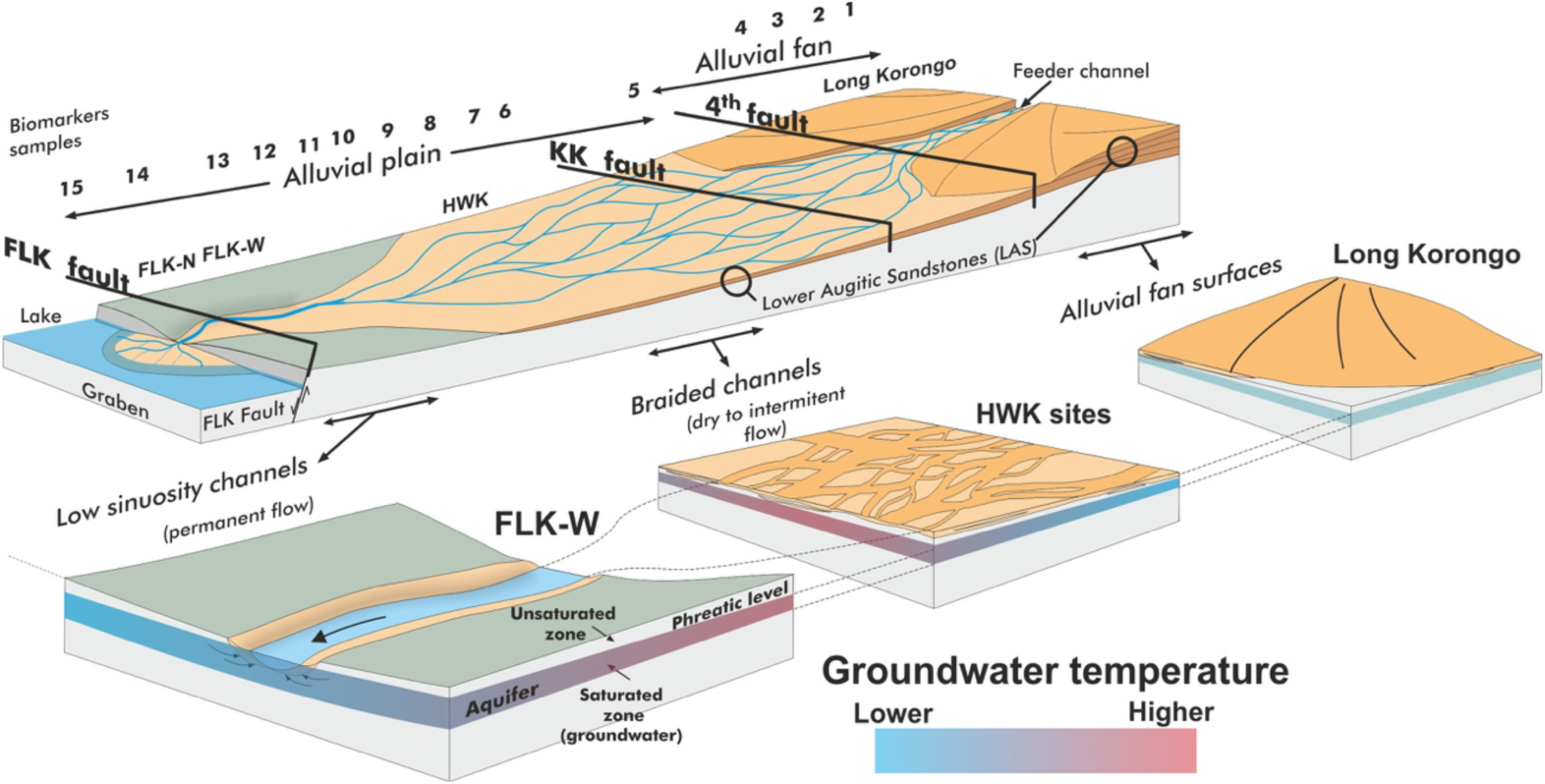
3D geomorphological reconstruction of the LAS unit between Long Korongo and FLK Fault during dry season.

Additional support for the presence of hyperthermophilic microbes comes from the *n*-alkanols and fatty acids, which are dominated by C_18_ chain-length in most of the samples, which has been also identified as a major component in other *Aquificales*^*46,47,52*^. Alcohols and fatty acids with longer chains (C_24_-C_30_) likely originate from both microbes and detrital plant organic matter (Fig.4C).

Isoprenoidal lipids identified in our study include cholesterol derived from animals, the phytosterols sitosterol, stigmasterol, campesterol, and stigmastanol, as well as amyrin, a plant triterpenoid. Interestingly, some samples contained 5β-stigmastanol, a known faecal biomarker for herbivore feces^57^. LAS 8 and 15 also showed an isoprenoidal profile dominated by archaeol, a methanogen biomarker^58^. LAS 5, 9, 10-13, and 15, also yielded a rare polycyclic triterpenoid, tetrahymanol, which is typically produced by ciliates and also by a few bacteria^59^ (SI2.C). Only a subset of methanotrophs and SRB seem to be able to synthesize tetrahymanol^59^, depending on the redox potential of the ecosystem. Tetrahymanol-producing aerobic methanotrophs are often identified in the suboxic zone of marine or freshwater bodies, while SRB occur in the anoxic sediments^59^.

Other microbial lipids have been also identified in the LAS samples. LAS 10-12 yielded significant amounts of fern-7-ene and fern-8-ene which have been attributed to ferns, but can also be produced in substantial amounts by bacteria^60^. Similarly, hop-17(21)-ene, hop-22(29)-ene which are prevalent in many bacteria^61^, were identified in some of the samples.

## LAS landscape and Hominin Evolution

Although the geothermal nature of Olduvai basin has not been studied in any detail, the presence of hydrothermal systems within the Tanzanian portion of the East African Rift are no surprise, as areas near Olduvai Gorge such as Lake Natron, Lake Manyara, Lake Eyasi, and Ngorongoro Caldera host hydrothermal activity today^41,62,63^. The deposition of the LAS sediments corresponds to an event with major fault activation, that, for the first time, enabled the reactivation and progradation of drainage networks to the Olduvai Basin, changing the nature of the sedimentation from lacustrine to fluvial^22^. Faults in the rift system are associated with deep-seated springs, characterized by the temperature and salinity of the water. Volcanism is also active at this time, and is characterized by the deposition of rich phenelinites-foidite tuffs and volcanic ashes whose pyrolysis products can be identified in the LAS sediments. The presence of hydrothermal waters does not appear exclusive to the LAS deposition event, and although no thermal biomolecular markers have been found yet during Bed I, diatom species specifically adapted to warm and hot waters, as well as illite/smectite layers potentially suggesting hydrothermalism^64^, have been previously identified before in the sequence^65^. The ubiquitous presence of silicified plants^36^ might also support hydrothermal activity, although this mineralisation can also be enhanced in other conditions.

Our biomarker evidence identifies a paleolandscape influenced by tectonics, volcanism, and hydrothermal activity which may have had important implications for early humans at Olduvai Gorge. Bailey and colleagues^12,66,67^ have suggested that tectonics and volcanism may have created advantages for hominin survival by creating dynamic landscapes that offered protection and resources. The particular tectonic landscape at Olduvai Gorge, fringed by major faults, offered a continuous supply of potable water from hot springs and rivers that would have attracted populations of large herbivores as attested by the fauna recovered in the archaeological sites^16^. The identification of 5β-stigmastanol in some of our samples, especially upstream and FLK-W, suggests a dominant input of herbivore faeces in these samples. Animals would have consumed the waters at Olduvai Gorge much as they do today in Yellowstone National Park, which would have provided a perfect opportunity for hominins to hunt large prey. The presence of biomarkers diagnostic for hyperthermophiles in some of our samples suggests that in parts of these streams or pools, the temperature reached as high as 80-90°C. At such temperatures, starches gelatinize, improving the accessibility of otherwise indigestible and toxic nutrients in roots and tubers, that became more available with the progressive loss of canopy in Bed II. Cooking animal tissues in high temperature waters would also have facilitated higher digestibility, palatability, and energy gain, with the benefit of adding nutrients derived from the hot spring water^68^. Wrangham and colleagues^21,68^ pointed to the evolutionary significance of cooking in human evolution, and suggested that cooking might have begun with *H. erectus*^68^.

While evidence of fire use is controversial during this time, hot springs may have offered *H. erectus* the opportunity to sustain the increased energetic costs of encephalization by providing an efficient way to thermally process the edible resources the wetland offered. Hominins may have sought these cooking opportunities given the preference that chimpanzees exhibit for cooked food and their willingness to transport raw food in anticipation for future chances of cooking^69^. Hydrothermal waters have traditionally been used for cooking in some areas of Japan, New Zealand, and Iceland, where the hot spring-cooked food is particularly appreciated for its nutrient content and tenderness.

## Conclusions

Elucidating the possible complex interactions between early humans and their environment at Olduvai Gorge provides context for understanding early hominin evolution. Current interpretations of the Olduvai landscape at 1.7 Myr ago rely on a ‘broad-strokes’ approach to environmental reconstruction, while hominins may have experienced different and more dynamic local conditions. Using this lens, we tested an initial hypothesis that differences between stone tool technologies at two sites in Olduvai Gorge, HWK and FLK-W, could be associated with specific local resources.

In this report, we provide evidence that the structural and isotopic composition of lipid biomarkers found in the 1.7 Myo LAS samples reveal a mosaic ecosystem, dominated by the presence of groundwater-fed rivers and aquatic plants. Further, we observe differences between the HWK and FLK-W sites. The former site appears drier, with intermittent water inputs, as suggested by the lower P_aq_ values, high pristane-to-phytane ratio, and TAR_FA_, indicative of higher terrestrial inputs. The low pristane-to-phytane ratio and the high P_aq_ of the latter together with the high P_alg_ (SI2.A), suggest a permanently waterlogged environment consistent with the wide channel previously described^14^. Phytolith evidence from FLK-W points to the presence of grasses and palms^70^, which may be supported by the predominance of the C_31_ *n*-alkane within the lipids from FLK-W. The palm shade likely supported the growth of *sphagnum* moss.

In the context of progressive aridification and expansion of savanna grasslands, the 1.7 Myo Olduvai Gorge paleolandscape depicted by our data was a groundwater-fed wetland with rivers partly sourced by hydrothermal waters and populated with patchy vegetation, dominated by C_3_ aquatic plants and angiosperm shrublands potentially including edible plants such *Typha*, sedges, ferns, and palms. This type of patchy vegetation is seen in African wetlands today^63^, and is highly dependent on water availability, temperature, and pH.

We also identify lipid biomarkers typical of hydrothermal features that are colonised by biofilm-forming microbes such as *Aquificales* and other thermophiles, with variations in community composition within the landscape. The interaction between tectonic dynamics (i.e., volcanism), hydrothermalism, and their influence on early human evolution has not been addressed before, except as theorized by Bailey and colleagues^12,66,67^. Here, we present the oldest geochemical evidence of a landscape-scale reconstruction associated with groundwater-fed habitats, in which hot springs and rivers partly sourced by thermal waters prompted intensive use of the landscape by hominins.

The accumulation of animal carcasses in the HWK and FLK-W sites suggest long-term, thorough hominin use of the space near hydrothermal features, which likely attracted large herbivores as suggested by our faecal biomarker record. Hot springs may have provided a convenient way to cook food that would have required minimal effort, which, at the same time, decreased digestibility-reducing and toxic compounds in starches. Cooking may, thus, have had a simple pre-fire stage during human evolution.

## Methods

### Sampling

Sampling was carried out in all stratigraphic sections described by Uribelarrea and colleagues^8.^ 6 additional outcrops, located between these sections, were also sampled in order to increase sample size for higher reproducibility and confidence. Throughout the studied area, the LAS were identified overlying the LD. Samples were collected from the basal 5cm of the LAS using sterilized tools, wrapped on combusted aluminium foil and stored in cloth bags.

### Biomarker extraction and isolation

Sediment samples were freeze-dried and powdered prior to extraction. After addition of recovery standard (1-pentadecanol), extraction was performed using and accelerator solvent extractor (Dionex ASE 300 system) with DCM and MeOH (9:1 v/v) using a method with 3 cycles of 15 min at 100°C and 1,500 psi. The total lipid extract was concentrated using a turbovap with a gentle stream of N2.

An aliquot of the total lipid extract (TLE) was derivatized by acid methanolysis (0.5M HCl in methanol) diluted in H_2_O and extracted with hexane:DCM (4:1 v/v). The concentrated derivatized extract was then separated into 3 fractions using a deactivated silica gel (2% H_2_O total weight) packed column by elution with hexane (F1), hexane:DCM (1:1) (F2) and DCM:MeOH (4:1). The polar fraction (F3) was then silylated using N,O-bis(trimethylsilyl)trifluoroacetamide (BSTFA).

### Biomarker analysis

Biomarkers were characterized by GC-MS with an Agilent 7890A series gas chromatograph interfaced with an Agilent 5977C mass selective detector. One microliter aliquots of apolar and derivatized extracts were injected in splitless mode onto a 60m fused-silica column (60m × 0.25 mm I.D. and film thickness of 0.25Um). The GC oven temperature was programmed as follows: 60°C injection and hold for 2 min, ramp at 6°C min to 300^a^C, followed by isothermal hold of 20 min. The MS was operated under conditions described above. Data was acquired, processed and identified under the conditions described above. 1-Pentadecanol was used as a recovery standard and internal injection and response factors were calculated for the different lipid classes using representative standards (aiC_22_, epiandrosterone, 1-nonadecanol and 2-methyloctadecanoic acid methyl ester). Procedural blanks were run in order to monitor background interferences.

The polar lipid fractions of LAS samples analysis were analysed by high performance liquid chromatography coupled with a quadrupole time of flight system (HPLC-QTOF). Analyses were conducted using the method established by Becker and colleagues in 2013. This method allows for the simultaneous analysis of long-chain alkenones, archaeal and bacterial glycerol dialkyl glycerol tetraether lipids (GDGTs), and long-chain diols. LAS polar lipid fractions were run on an Agilent Technologies 1260 Series HPLC coupled to an Agilent Technologies 6520 Accurate-Mass Q-TOF. 2% aliquots of the LAS TLE were dissolved in *n*-hexane:propan-2-ol (99.5:0.5, v:v) and were injected onto coupled Acquity UPLC BEH Amide 2.1 × 150mm × 1.7um columns (Waters), each maintained at 50°C. Separation was achieved by a method established by Becker et al., 2013. All LAS samples were run with a 10ng spike of an instrumental standard, a C46 glycerol trialkyl glycerol tetraether lipid (C46-GTGT). Procedural blanks were run alongside the LAS samples on the HPLC-QTOF to monitor instrument background as well as to detect possible contamination. MS^2^ spectra were also collected in data dependent mode, with three MS^2^ experiments targeting abundant ions with N_2_ as a collision gas.

Measurements were performed in full-scan mode, scanning m/z 100 to 2000 at 2 spectra/second. The machine was calibrated before running samples, and the typical error of mass accuracy was <1 ppm. Peak areas were corrected to the C46-GTGT standard before analysis. Analyses of the chromatograms from all instruments were conducted using Agilent MassHunter Qualitative Analysis Navigator version B.08.00. All peaks were identified based on their exact masses, corresponding MS2 spectra, and their retention times.

### Compound-specific isotopic analyses

Apolar fractions were subsequently analysed to characterize their isotopic signatures using a Trace 1310 interfaced with a GC Isolink II connected to a Thermo MAT 253. One microliter aliquot was injected in splitless mode onto a 60m DB5 column (60m × 0.25 mm I.D. and film thickness of 0.25Um) before combustion over cooper, nickel and platinum wire with oxygen and helium at 1000°C. Isotopic values were normalized using a mixture of n-alkanes (C_16_-C_30_) of known isotopic composition (mixture A5 of Dr. Arndt Schimmelmann). The standard deviation for the instrument, based on replicate standard injection was calculated to be below 0.4‰. All measurements were performed in duplicate. We only report here well-resolved analytes, corresponding to major compounds within the lipid classes.

### Bulk geochemical analyses

For bulk analysis, about 3g of sediment, previously freeze-dried and ground, was reacted with 1N HCl to remove inorganic carbon, then rinsed until reaching neutral pH. Analyses were carried on via elemental analysis-isotope ratio mass spectrometry (EA-IRMS) for organic carbon (TOC), total nitrogen (TN) and carbon isotopic composition of bulk organic matter (δ^13^C_TOC_) using an ECS 4010 from Costech interfaced to a Thermo Finnigan Delta Plus XP.

## Acknowledgements

We thank the Tanzania Commission for Science and Technology, the Department of Antiquities and the Ngorongoro Conservation Area Authority for field permits and support. Archaeological research at Olduvai Gorge is funded by the Spanish Government I+D project (HAR2017-82463-C4-1-P). Research at MIT was supported by a Marie Skłodowska Curie Global Fellowship (H2020-MSCA-IF-2016-750860) to A.S., and a grant (NNA13AA90A) from the NASA Astrobiology Institute to R.E.S. We are grateful to the crew at the TOPPP archaeological project in Olduvai. We also thank X. Zhang and G. Izon for their assistance with compound-specific isotope analyses, and T. Evans for his assistance with the HPLC Q-TOF.

