## Supplementary Information for "The role of tectonics and hydrothermalism in early human evolution at Olduvai Gorge"

### 1- Geological setting additional information

Geologically, the Olduvai basin lies between the mountains of metamorphic rocks to the north and large volcanoes to the south. Throughout the Quaternary it has been filled with 100 m of lake, river, alluvial, volcanic and wind sediments. This record of 100 m thick was divided by Hay (1976) into 7 large units: Bed I, II, III, IV, Masek, Ndutu and Naisiusiu (Fig.1). The entire basin is crossed by 7 major faults, associated with the great African East Rift, 1st, 2nd, 3rd, 4th, KK, FLK and 5th faults (Fig.2). The nearest volcanoes are the Lemagrut (~ 15km) and the Ngorongoro (~ 20km).

During Bed II (1.75-1.35 m.y.) the paleogeography was dominated by a large central lake, fed by the rivers that came from the slopes of the volcanoes and the metamorphic reliefs to the north. In the study area, the beginning of Bed II is represented by lake margin facies, several meters of waxy claystone deposited in a very quiet environment. Interbedded, the aeolian-volcanic complex "Lemuta member" was formed, which contains the marker tuff IIA (1.70-1.75 m.y.). The tectonic activity in Olduvai begins precisely during Bed II. A graben is formed between faults FLK and 5th. There is a decrease in the base level that reactivates the entire drainage network in the region. As a consequence, an important erosion of the elevated blocks takes place and a great paleosurface develops, representing the Lower Disconformity (LD). Probably, the same fluvial channels that formed this disagreement deposited great amounts of augitic volcanic sand, that has been called Lower Augitic Sandstone (LAS). It is an entire fluvial system, represented, upstream downstream, by large alluvial fans (Long Korongo). Towards the NW, LAS is represented by a system of small channels of braided type with mixed and coarse bedload (HWK area). These channels are concentrated downstream, forming isolated sinuous channels in an alluvial plain (FLK-W), which connects to the edge of the central lake. These channels are deeper and narrower than those upstream. Thus, during periods of water scarcity, the channel in FLK-W is maintained with water, while the braided channels are dry[1,2].

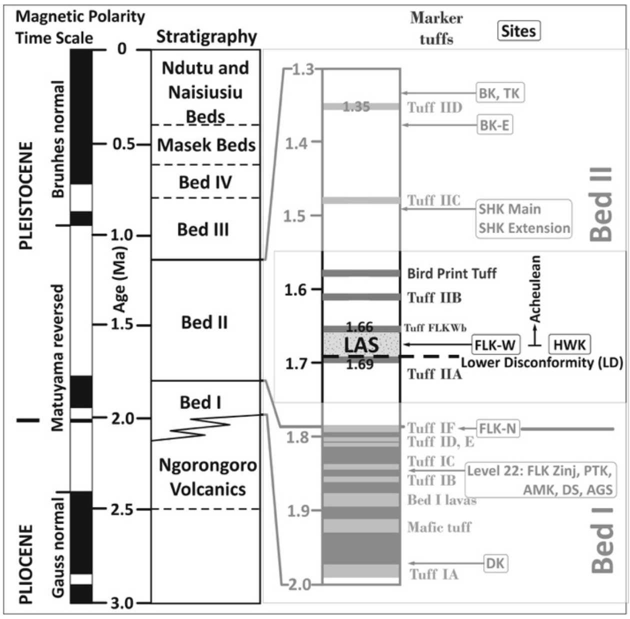


Figure 1. Composite stratigraphic section for Olduvai Gorge, showing the relative order of the 7 geological units. The Bed I and Bed II have been highlighted and within this last one, the appearance of the oldest Acheulean of the Gorge just above the Lower Disconformity (LD) (dashed line), in the Lower Augitic Sandstones (LAS).

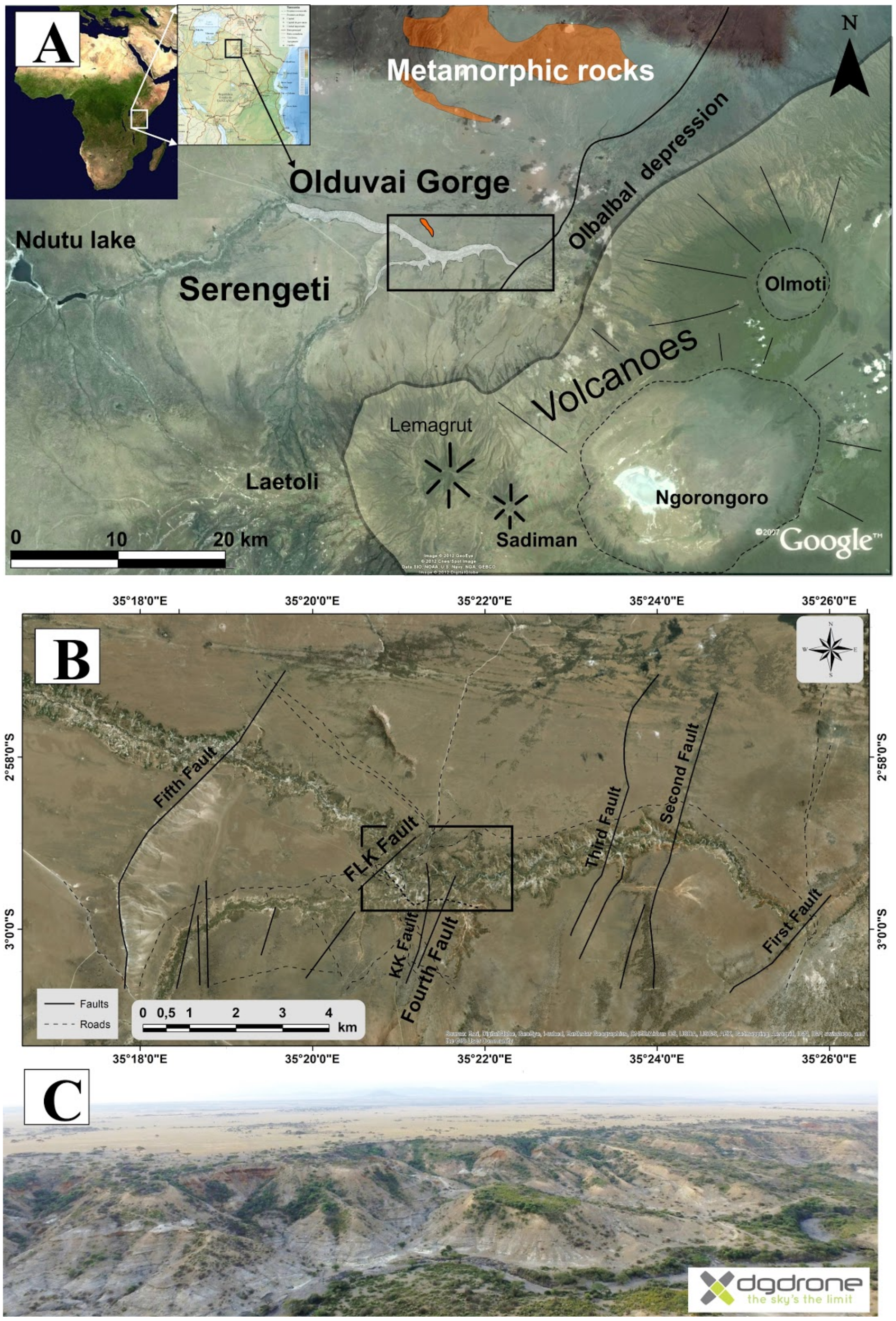


Figure 2. A, General view of the south of the Serengeti, in the North of Tanzania. To the southeast are the great volcanoes Lemagrut, Sadiman, Ngorongoro and Olmoti. To the north the mountain ranges of metamorphic rocks (orange). B, Olduvai Gorge. The main faults have been labelled. C, oblique aerial photograph taken from an unmanned aerial vehicle, of the main gorge.

### 2- Lipid Biomarkers additional information

A. Table with lipid proxies

| *sample* | $\delta^{13}C_{31}$ | $\delta^{13}C_{25}$ | $P_{aq}$ | $C_{33}/C_{31}$ | $TAR^{FA}$ | $P_{alg}$ | *TOC* | $\delta^{13}C_{TOC}$ | *Pr/Ph* | *ACL* | *CPI* |
|---|---|---|---|---|---|---|---|---|---|---|---|
| LAS 1 | -24,94 | -24,94 | 0,23 | 0,66 | 0,01 | 0,04 | 0,02 | -25,37 | 1,81 | 29,33 | 2,24 |
| LAS 2 | -24,69 | -28,32 | 0,31 | 0,92 | 0,01 | 0,04 | 0,02 | -27,35 | 0,62 | 30,71 | 3,15 |
| LAS 3 | -25,41 | -29,16 | 0,21 | 0,73 | 0,05 | 0,32 | 0,04 | -24,65 | 0,43 | 29,21 | 3,38 |
| LAS 4 | -22,85 | -19,92 | 0,57 | 0,56 | 0,03 | 0,11 | 0,03 | -21,82 | 0,78 | 27,40 | 4,22 |
| LAS 5 | -23,59 | -22,64 | 0,30 | 0,64 | 0,03 | 0,02 | 0,02 | -23,96 | 0,00 | 28,73 | 4,35 |
| LAS 6 | -23,81 | -24,60 | 0,16 | 0,68 | 0,22 | 0,00 | 0,03 | -22,40 | 1,95 | 29,68 | 4,14 |
| LAS 7 | -26,75 | -30,34 | 0,38 | 0,63 | 0,03 | 0,01 | 0,07 | -28,85 | 0,11 | 28,56 | 1,51 |
| LAS 8 | -25,42 | -27,81 | 0,00 | 0,59 | 0,00 | 0,43 | 0,07 | -28,80 | 0,20 | 0,00 | 0,00 |
| LAS 9 | -24,57 | -25,76 | 0,38 | 0,48 | 0,03 | 0,95 | 0,19 | -23,20 | 1,02 | 27,92 | 2,28 |
| LAS 10 | -25,11 | -27,08 | 0,38 | 0,43 | 0,02 | 0,27 | 0,15 | -24,99 | 0,67 | 28,15 | 2,16 |
| LAS 11 | -23,85 | -25,70 | 0,33 | 0,48 | 0,02 | 0,04 | 0,08 | -27,10 | 0,36 | 28,61 | 2,72 |
| LAS 12 | -27,02 | -29,20 | 0,40 | 0,42 | 0,02 | 0,03 | 0,09 | -26,32 | 0,51 | 28,14 | 1,29 |
| LAS 13 | -25,41 | -28,56 | 0,36 | 0,40 | 0,01 | 0,05 | 0,13 | -26,20 | 0,22 | 28,08 | 1,51 |
| LAS 14 | -22,95 | -25,24 | 0,29 | 0,96 | 0,14 | 0,75 | 0,03 | -27,77 | 0,36 | 29,99 | 3,99 |
| LAS 15 | -23,80 | -26,45 | 0,45 | 0,00 | 0,02 | 0,00 | 0,09 | -27,73 | 0,00 | 27,78 | 2,28 |

Table 1. Various organic geochemical proxies for organic matter preserved in outcrops from Eastern fluvial lacustrine deposits (eastern lake-margin) dated up to 1.7 Ma at Olduvai Gorge. **$\delta^{13}C_{31}$**: Plant lipid $\delta^{13}C$ values for $n_{31}$ alkane (higher values, more $C_4$ vegetation). **$\delta^{13}C_{25}$**: Plant lipid $\delta^{13}C$ values for $nC_{25}$ macrophytic alkane (higher values, more macrophytes). **$P_{aq}$**: Ratios of macrophytic lipids ($nC_{23}$ + $nC_{25}$) relative to macrophytic and terrestrial lipids ($nC_{23}$ + $nC_{25}$ + $nC_{29}$ + $nC_{31}$). **$C_{33}/C_{31}$**: Ratio of $nC_{33}$ to $nC_{31}$, (higher values, more grasses) **$TAR_{FA}$**: Terrigenous to aquatic n-alkanoic acids ratio reflecting the importance of terrigenous and aquatic sources $(C_{24} + C_{26} + C_{28})/(C_{14} + C_{16} + C_{18})$ (higher values, more terrestrial input). **$P_{alg}$**: Ratios of algal lipids ($nC_{17}$ + $nC_{19}$) relative to algal and terrestrial plant lipids ($nC_{17}$ + $nC_{19}$ + $nC_{29}$ + $nC_{31}$) higher values, more algal input). **TOC**: Total organic carbon, weight percent (higher values, more organic matter production). **$\delta^{13}C_{TOC}$**: Carbon isotopic composition of total organic carbon (higher values, more C4 vegetation). **Pr/Ph**: Ratios of pristane to phytane (values above 2 often indicative of more terrestrial plant input, values below 1 indicative of anoxic conditions, values around 1 indicative of oxic/anoxic alternating conditions). **ACL**= $\sum(C_n \times n) / \sum(C_n)$ Average chain length of individual n-alkane abundances. **CPI**= $[\sum_{odd}(C_{21-33}) + \sum_{odd}(C_{23-35})]/(2\sum_{even}C_{22-34})$ Carbon preference index, indicative of the abundance of odd over even carbon chain lengths (lower CPIs often indicative of microbial degradation or maturation of the sample).

B. MAGEs Identification

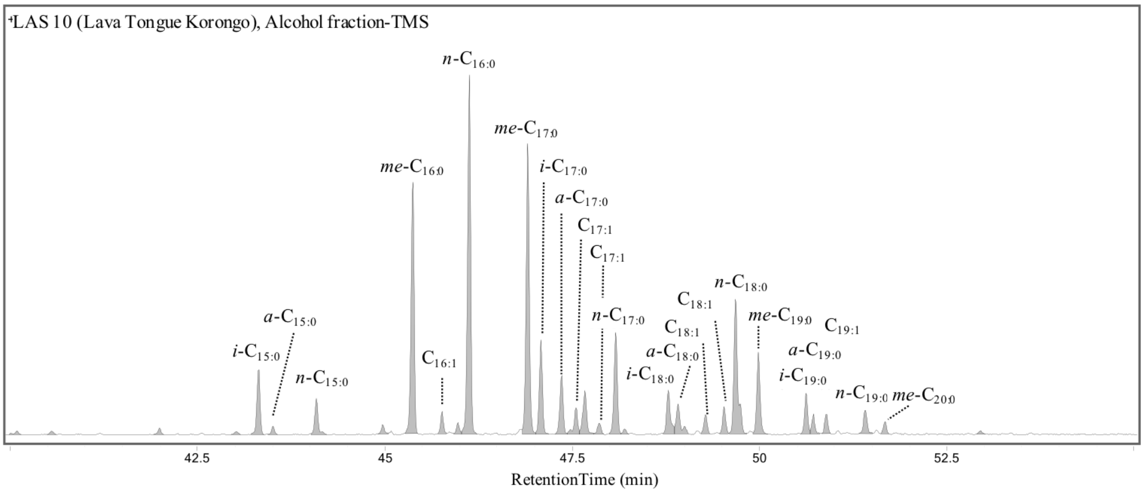


Figure 3. Partial mass chromatogram of the alcohol fraction extracted from sample LAS 10 of the diagnostic fragment of MAGEs (*m/z* 205). Peaks are labelled with the carbon number of attached alkyl moieties and the number of double bonds in the alkyl chain.

### C. Tetrahymanol Identification

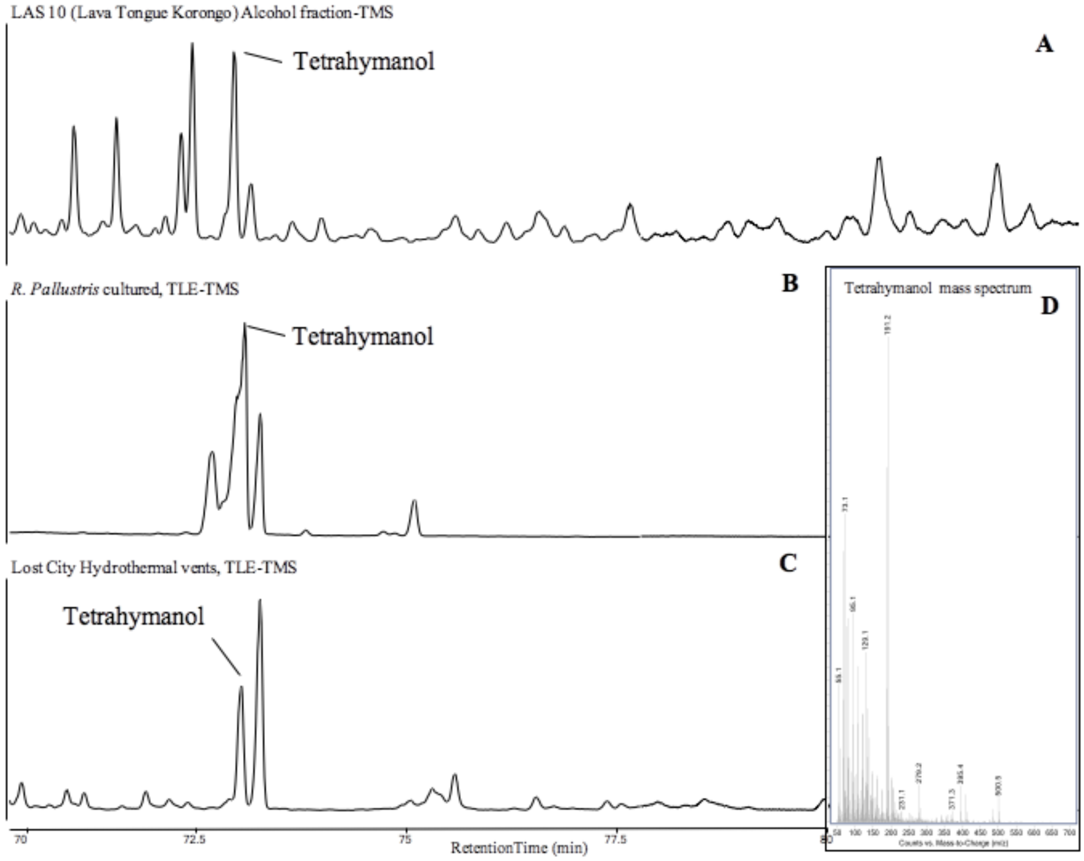


Figure 4. Partial mass chromatogram of tetrahymanol extracted from A) the alcohol fraction of LAS 10, B) *Rhodopseudomonas pallustris* culture, and C) a sample from Lost City hydrothermal vents[3], D) Mass spectrum of tetrahymanol extracted from the alcohol fraction of LAS 10.

D. *T. ruber* fatty acids

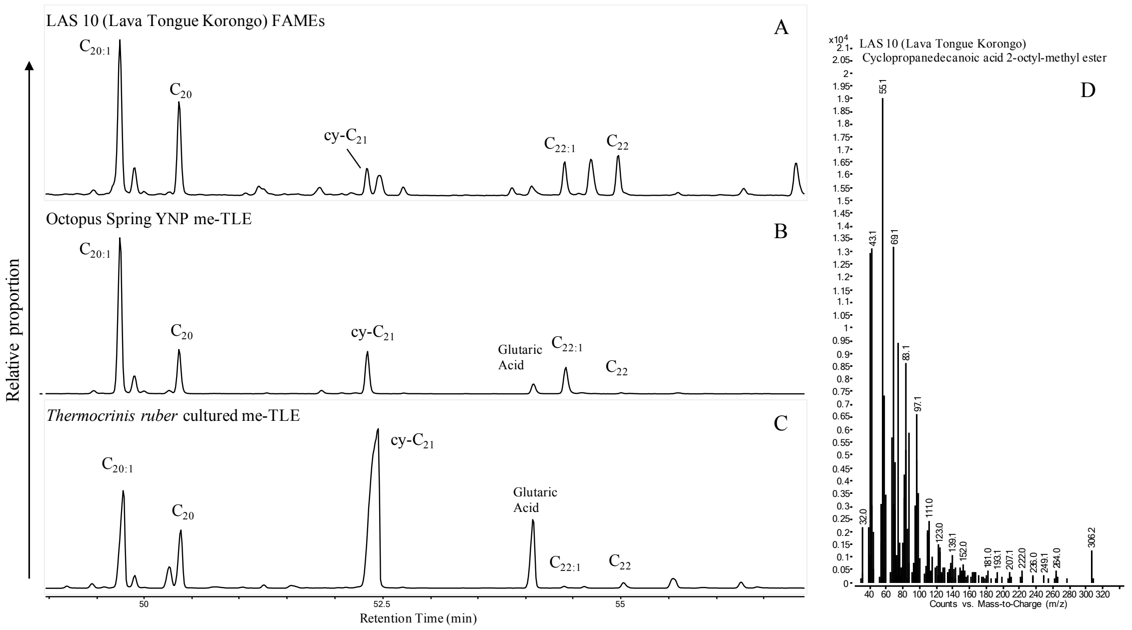


Figure 5. Partial mass chromatogram of fatty acids methyl esters showing the presence of $C_{20:1}$, $C_{22:1}$ and cy-$C_{21}$ extracted from A) the FAMEs of LAS 10, B) Octopus Spring sample from Yellowstone National Park[4,5], and C) a sample cultured of *Thermocrinis Ruber*, D) Mass spectrum of Cy-$C_{21}$ fatty acid extracted from LAS 10.

E. GDGT analyses

Glycerol dialkyl glycerol tetraethers (GDGTs) are ether-linked membrane lipids produced by both archaea and bacteria. Isoprenoidal and branched GDGTs, produced by archaea and bacteria respectively, are often methylated and can contain cyclopentyl moieties[6–8]. Structures of GDGTs may be linked to ambient environmental conditions at the time of production[6,9–11]. Two main indices are used to evaluate the extent of methylation and cyclization in brGDGTs—the methylation of branched tetraethers (MBT) and the cyclization of branched tetraethers (CBT)[12]. The MBT and CBT indices have been calibrated to reconstruct air temperatures, and GDGT distributions in globally-distributed soils have been found to correlate significantly with mean air temperatures (MAT), with the degree of methylation corresponding to temperature and the degree of cyclization correlating with pH[12].

While both bacterial and archaeal GDGTs were detected in the Olduvai samples, their abundances were too low for reliable quantification in possible paleoclimate reconstructions. Ferland[13] examined GDGTs in the paleolake Magadi, and found similar absences and low abundances of GDGTs in their lacustrine sediments.

When run on the HPLC-QTOF using the method established by Becker and colleagues in 2013[14] and quantified with regards to an instrumental C46-GTGT standard[15], few complete suites of known bacterial GDGT lipids were detected within the LAS Samples. Only one out of sixteen samples, LAS 4, contained the entire brGDGT suite necessary to calculate the MBT and CBT ratios used in paleotemperature reconstruction, established by Weijers and colleagues in 2007. Using ratios of compound abundances detected by

HPLC-QTOF, we were able to estimate the mean air temperature (MAT) for LAS 4 using the MBT' ratio established by Peterse and colleagues in 2012[16].

MBT' = (Ia + Ib + Ic) / (Ia + Ib + Ic + IIa + IIb + IIc + IIIa)
CBT = -log ((Ib + IIb)/(Ia + IIa))

By substituting the peak concentrations adjusted to the instrumental C46-GDGT standard, we were able to glean the following temperatures from the LAS 4 polar fraction using the following four air temperature calibrations.

| Calibration | Weijers et al., 2007[12] | Tierney et al., 2010[11] | Sun et al., 2011[17] | Loomis et al., 2012[18] |
|---|---|---|---|---|
| T (°C) | 1.841062123 | 16.59080891 | 12.67466515 | 10.17604657 |

Tierney et al.'s 2010 calibration aligns most closely with the suggested temperatures for Bed I and Bed II times (16ºC), which were about 5 to 7ºC lower than today's mean annual temperature (22ºC) at Olduvai Gorge[19,20].

3- Implications for human evolution additional discussion

The interaction between tectonic dynamics (i.e., volcanism) and hot spring environments and their influence on human evolution has not been considered from a perspective of natural selection. Here, we have documented an association between thermal hot springs and rivers and the archaeological record contained in the LAS paleolandscape. All the archaeological sites associated with this paleoecosystem are deep vertical deposits (the HWK complex) or sequences of multiple archaeological levels (i.e., FLK-W) indicating a redundant and prolonged use of the space by hominins and other mammals. The taphonomic study of the LAS sites indicate that they were palimpsests with a high input by carnivores[21,22]. Only FLK-W contains a higher anthropogenic impact[23]. The intensive use of the space by hominins and other mammals is evidenced by the continuous accumulation of remains (fossil bones and stone tools) over a vast area covering the braided-river system comprised between HWK and HWKEE[24,25] with a decreasing density in the Long Korongo and towards the north. It is precisely in this HWK complex area that the highest concentration of herbivore faecal biomarkers has also been found.

Evidence of carcass butchery and consumption in the HWK complex area, within the braided river system habitat, is rather scarce; very few percussed and cut-marked bones (37 at HWKEE and 6 at HWK 1-2) have been reported from such a section of the landscape[21,26]. In contrast, the density of stone tools is astounding. And most surprisingly, they do not seem to be discreetly clustered around focal points, as is documented in most Bed I sites, but they are widespread over an extensive area covering thousands of square meters around the HWK site complex. Once we move from the HWKEE-HWKE-HWK-RHS1 paleolandscape, both towards the FLK-W area in the north and the Long Korongo in the south, both fossil bones and stone artefacts decrease drastically and are found only sporadically in the form of low-density scatters. The HWK complex also includes a moderate density of remains modified by carnivores, including felid-modified carcasses as well as a conspicuous intervention by hyenas[21,22]. The overlap in the use of this space by fissiped carnivores and hominins over vast amounts of time created a large-scale palimpsest in the form of vertically dense deposit. The HWK area was, therefore, a highly productive paleolandscape; high depositional rates of bones and stone tools created this rich archaeological area. Although the presence of fossils and stone tools seems continuous over this rich area, this appreciation needs to be confirmed with a landscape archaeology project aiming at creating regular sampling trenches in the spaces in between sites.

Given that the evidence of hominin exploitation of carcasses in this area is so scant, the large amount of battering Oldowan tools found could probably be related to the exploitation of plant resources. Most of the discrete evidence of processing of carcasses by hominins on that paleolandscape is found at the Acheulian site of FLK-W[27]. A grassland and marshland dominated the braided-river environment and palm trees have been found at FLK-W[28], where the river morphed from a braided system to a wide channel before entering the lake (Uribelarrea et al., 2017). The lithic assemblage of the HWK complex has been typologically and technologically classified as Oldowan[24,25]. The presence of Acheulian industries has been typologically and technologically documented at FLK-W[27,29]. This indicates a clear co-existence of both technologies on the same environment. This also emphasizes the need to adopt an adaptive approach to understand lithic variability across paleolandscapes.

Future research should test if the evidence of intensive use of the space by hominins associated to habitats where thermalism is prominent is a coincidence or not. In the latter case, there is an important adaptive component of these environments in the evolution of human behaviour that must be properly understood. A hypothesis to be tested is that such habitats could have been preferred because hominins may have easily engaged into cognitively-simple cooking behaviours prior to the use of fire.

### References

1. Uribelarrea, D., Martín-Perea, D., Diez-Martín, F., Dominguez-Rodrigo, M., Baquedano, E., Mabulla, A. A geoarchaeological reassessment of the co-occurrence of the oldest Acheulian and Oldowan in a fluvial ecotone from lower middle Bed II (1.7ma) at Olduvai Gorge (Tanzania). Quaternary International
